# Oxidation state dependent conformational changes of HMGB1 regulate the formation of the CXCL12/HMGB1 heterocomplex

**DOI:** 10.1101/555946

**Authors:** Enrico M. A. Fassi, Jacopo Sgrignani, Gianluca D’Agostino, Valentina Cecchinato, Maura Garofalo, Giovanni Grazioso, Mariagrazia Uguccioni, Andrea Cavalli

## Abstract

High-mobility Group Box 1 (HMGB1) is an abundant protein present in all mammalian cells and involved in several processes. During inflammation or tissue damage, HMGB1 is released in the extracellular space and, depending on its redox state, can form a heterocomplex with CXCL12. The heterocomplex acts exclusively via the chemokine receptor CXCR4 enhancing leukocyte recruitment.

Here, we used multi-microsecond molecular dynamics (MD) simulations to elucidate the effect of the disulfide bond on the structure and dynamics of HMGB1.

The results of the MD simulations show that the presence or lack of the disulfide bond between Cys23 and Cys45 modulates the conformational space explored by HMGB1, making the reduced protein more suitable to form a complex with CXCL12.

## 1. Introduction

High-mobility Group Box 1 (HMGB1) is an abundant chromatin-associated protein present in all mammalian cells. It is formed by 215 amino acids, divided into two domains, “BoxA” (Gly2-Ile79) and “BoxB” (Phe89-Arg163), connected by a nine amino acid loop, and a highly disordered negatively charged C-terminal tail.

BoxA contains a pair of cysteines (Cys23 and Cys45) that can form a disulfide bond under oxidative conditions. In contrast, only one unpaired cysteine is present in BoxB (Cys106, Figure 1A) [1, 2].

**Figure 1.**
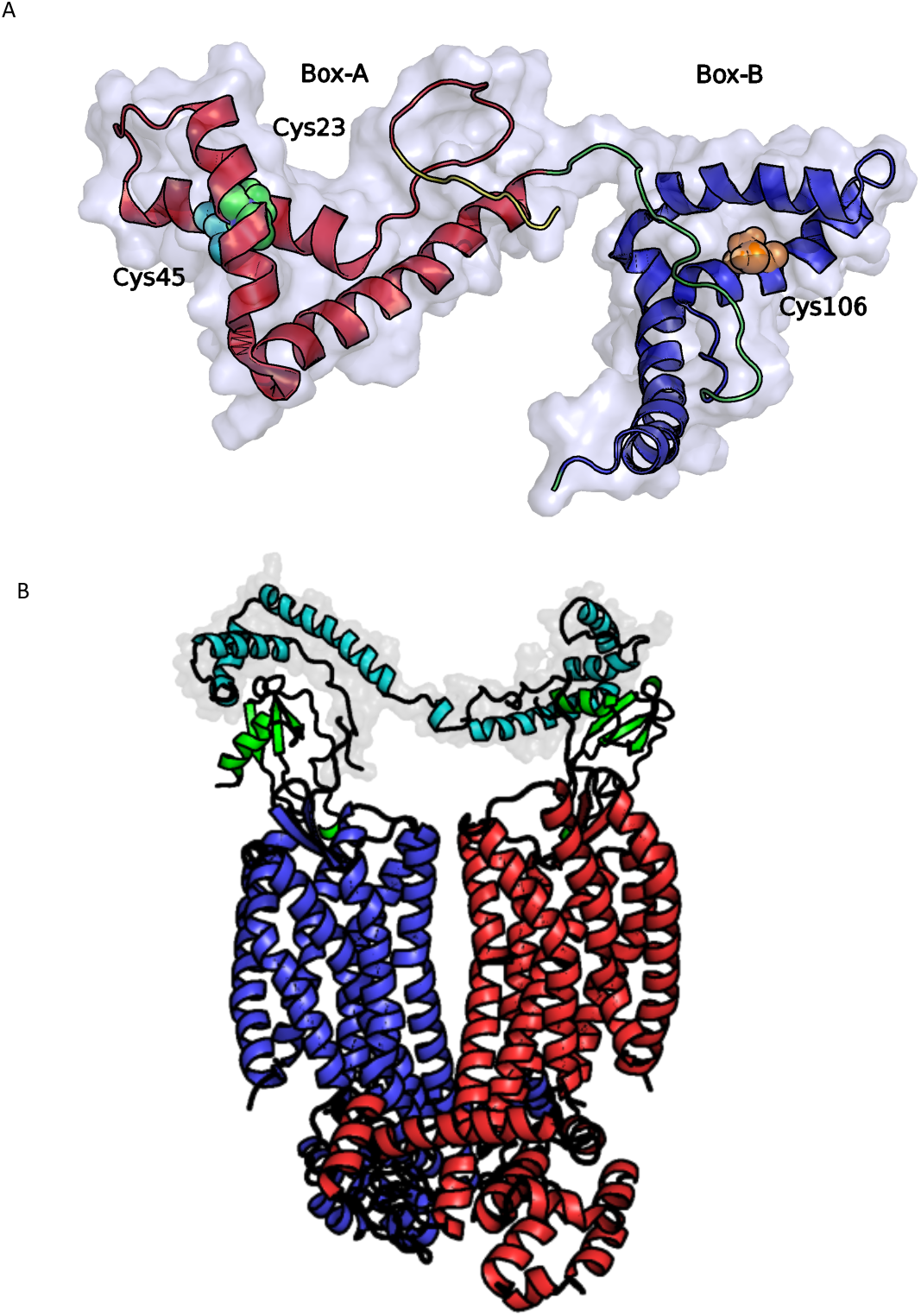
(A) Structure of HMGB1 (PDB ID code 2YRQ) solved by NMR. Protein domains are presented in different colors: BoxA (red), BoxB (blue), and the loop between the two domains (green). The three cysteines located at positions 23 and 45 in BoxA and 106 in BoxB are displayed as van der Waals balls in different colors. (B) Explicative representation, of the binding of the CXCL12_2_/HMGB1 to a CXCR4 dimer. HMGB1 is depicted in aquamarine, the two CXCL12 in green, while the two CXCR4 monomers in blue and red respectively.

The three domains of HMGB1 play a key role in establishing and regulating its wide interactome [3, 4], as well as, in the modulation of the protein conformation [5].

Depending on its cellular localization, HMGB1 performs different functions. In fact, as a nuclear protein, it is involved in DNA repair, transcription, telomere maintenance, and genome stability [2, 6, 7], while during cellular death or inflammation, HMGB1 is released in the extracellular space where it functions as an alarmin [8, 9].

According to multiple studies, several HMGB1 functions depend on its redox states [10, 11].

The nuclear and cytosolic environments are characterized by a negative redox potential that maintains HMGB1 in reduced form (fr-HMGB1). During an inflammatory process, the extracellular space, enriched in reactive oxygen species, lead to the formation of a disulfide bond between cysteines at positions 23 and 45 of BoxA (ds-HMGB1) [12]. ds-HMGB1 activates Toll-like Receptor 2 (TLR2) and 4 (TLR4) inducing the release of proinflammatory chemokines and cytokines activating innate and adaptive immune responses. On the contrary, fr-HMGB1 binds to the receptor for advanced glycation end products (RAGE), modulating autophagy [9, 13, 14].

The CXC ligand 12 (CXCL12) is expressed in many tissues both under homeostatic and inflammatory conditions and can stimulate cellular recruitment by activating the CXC chemokine receptor type 4 (CXCR4) [15]. In 2012, researchers in our group have shown that the CXCL12/HMGB1 heterocomplex enhanced the activities ofCXCR4 in human monocytes [16]. In particular, a suboptimal concentration of CXCL12, which per se would not trigger any chemotactic response, efficiently promotes migration of human monocytes, by forming a heterocomplex with fr-HMGB1 [16, 17]. More recently, other studies demonstrated the important role of the heterocomplex in tissue regeneration [13, 18, 19] and in fueling the inflammatory response in patients with Rheumatoid Arthritis [20].

A particular feature of the CXCL12/HMGB1 heterocomplex is that only fr-HMGB1 can complex with CXCL12, promoting CXCR4-induced response [17]. This appears contradictory because the extracellular space, where the heterocomplex is formed, is rich in reactive oxidative species [17]. However, under specific conditions, cells can release glutathione reductase and enzymes of the thioredoxin system to counteract the oxidative stress in the microenvironment, contributing to maintain HMGB1 in the reduced state [20, 21].

While a structure of the heterocomplex is currently unavailable, NMR chemical shift mapping clearly showed an interaction between CXCL12 and the two domains of HMGB1 (BoxA and BoxB), separately [16]. Furthermore, the same experiments showed that the binding of CXCL12 to HMGB1 induces conformational changes in the *N-*terminal domain of CXCL12 which is required to trigger the activation of the receptor. Based on these data, it was hypothesized that the heterocomplex is formed by two CXCL12 molecules bound to fr-HMGB1 (one to BoxA and one to BoxB), and that it would bind CXCR4 dimers (Figure 1B) [16].

In this study, aiming to validate the assumed mode of action of the heterocomplex, we applied several molecular modeling techniques, such as molecular dynamics (MD) simulations and protein-protein docking, to investigate which structural and/or conformational differences between the two redox states of HMGB1 could explain the different affinity of fr- and ds-HMGB1 for CXCL12.

According to our findings, ds-HMGB1 tends to be more compact and displays a lower accessible surface than fr-HMGB1, while the structure of BoxA remains essentially unchanged in the two states. Furthermore, in-depth analysis of the simulations and the results of protein-protein docking calculations showed that the vast majority of the conformations assumed by fr-HMGB1 are able to bind two CXCL12 molecules with an orientation and distance optimal to trigger the activation of CXCR4 dimers. We, therefore, propose that functional differences between fr- and ds-HMGB1 are at least partially caused by global changes in the configurational landscape of HMGB1.

## 2. Methods

The affinity of the CXCL12/HMGB1 heterocomplex was measured by microscale thermophoresis (MST) [22, 23]. Briefly, 100 nM HMGB1-His tagged, either reduced or oxidized, was labelled with 100 μM Monolith NTTM His-Tag Labeling kit RED-tris-NTA (L008, NANOTEMPER, Munich, Germany) 30 min at room temperature (RT) in the dark, and centrifuged (14.000 rpm; 10 min; 4°C) to discard the excess of dye in the tube. Labelled HMGB1 was used at final concentration of 10 nM in the presence of different doses of CXCL12, prepared performing 16 serial dilutions from the initial concentration of 14 μM according to the manufacturer instructions. Dilution buffer was obtained by mixing 20 mM NaCl/ NaH_2_PO_4_ pH 6.0 and PBS 0.1% Tween 20 pH 7.4 at 1:1 ratio.

Measurements were performed using the Monolith NT.115 MST Premium Coated Capillaries (K005, NANOTEMPER, Munich, Germany), excitation Power 20%, MST power medium, with the Monolith NT.115 Pico instrument (NANOTEMPER, Munich, Germany).

Apparent *K*_d_ values were computed fitting the compound concentration-dependent changes in normalized fluorescence (Fnorm) by the MO Affinity Analysis software provided by Nanotemper. Final results were obtained averaging four independent experiments.

Both CXCL12 and HMGB1 were prepared as in ref. [20]. Oxidized HMGB1 was obtained after sample dialysis, to remove DTT, and incubating the protein over night at room temperature to allow spontaneous oxidation.

### 2.2 Systems setup and MD simulations

MD simulations are powerful tools already applied to the study of some mechanistic aspects of the HMGB1 cellular functions [24, 25].

In this case, the HMGB1 structure solved by NMR spectroscopy (PDB ID 2YRQ), was used as a starting point for the simulations. As the first residue (Met1) of the protein is cleaved during posttranscriptional processing [26], this amino acid was deleted from the model and only the region from Gly2 to Arg170 (i.e., BoxA, BoxB, and the connecting loop) was considered in the MD simulations.

All the investigated HMGB1 models (fr- of ds-) were first minimized using the program ALMOST [27]. Then, the TLEAP module of AmberTools16 was used to solvate the protein in a box of water with a minimum distance of 10 Å from the protein surface. The net charge of the system was neutralized by adding a proper number of ions (17 or 15 Cl^-^ for fr- or ds-HMGB1 respectively). The ff14SB [28] force field parameters were used to describe the protein, while the TIP3P [29] model and the parameters proposed by Joung et al. [30] were used for water and counter ions, respectively. The solvated system was relaxed by a two-step protocol to remove atomic clashes [31] First, we performed an energy minimization for 10,000 steps, or until the energy gradient of 0.2 kcal/mol/Å was reached, restraining the atomic coordinates of backbone with harmonic potential (k=20 kcal/mol/Å^2^). This first phase was followed by an energy minimization for 100,000 steps or until an energy gradient of 0.0001 kcal/mol/Å was reached, without any restraint. After minimization, the temperature of the system was gradually increased to 300 K over 40 ps under constant volume condition (NVT) constraining the backbone coordinates in the first 20 with a harmonic potential (k=20 kcal/mol/Å^2^). Finally, the system was equilibrated at 300 K for 20 ps under constant pressure conditions (NPT, 1 atm). Pressure and temperature were maintained constant using the Berendsen barostat and thermostat, respectively [32]. Electrostatic interactions were treated with PME[33] with a cutoff of 9 Å. During the calculations, all bonds involving hydrogen atoms were constrained with the SHAKE [34] algorithm. All calculations were performed using the PMEMD of Amber16 code in the GPU accelerated version [35] with a time step of 2 fs.

Production runs were carried out using the following scheme. After the first simulation of 1 *µ*s, 29 of the saved frames were randomly selected and used as a starting point for 29 additional simulations (see Table 1). The atom velocities were reassigned at the beginning of each simulation to obtain uncorrelated and independent trajectories.

**Table 1.** Summary of the MD simulations performed in this study.

| System | Description | Simulation Time |
| --- | --- | --- |
| fr-HMGB1 | HMGB1 NMR structure (PDB ID code 2YRQ) | 30x1 μs |
| ds-HMGB1 | HMGB1 with a disulfide bond between Cys23-Cys45 | 30x1 μs |
| fr-HMGB1(I) | First representative cluster of the fr-HMGB1 + CXCL12<sub>2</sub> complex | 3x500 ns |
| fr-HMGB1(II) | The second representative cluster of the fr-HMGB1 + CXCL12<sub>2</sub> complex | 3x500 ns |
| ds-HMGB1(II) | The second representative cluster of the ds-HMGB1 + CXCL12<sub>2</sub> complex | 3x500 ns |

### 2.3 Trajectory Analysis

HMGB1 radius of gyration (RoG) was computed using the cpptraj [36] module available in AmberTools16 including all the protein residues. To assess the convergence of RoG calculation, 75000 snapshots sampled over the 30×1 *µ*s trajectories were divided into six groups of 12500 snapshot. Then the snapshots belonging to one of the six groups were excluded from the calculation and the results compared with those obtained using the full conformation ensemble (Figure S2).

The RMSDs of BoxA (Lys8 to Ile79) and BoxB (Lys96 to Arg163) were computed with the VMD [37] software, using the first conformation from the HMGB1 NMR bundle (PDB ID code 2YRQ) as a reference.

The solvent accessible surface area (SASA) was computed for the entire protein, BoxA, and BoxB using the LCPO algorithm [38] implemented in the cpptraj module of Amber16. Finally, atom-atom and residue-residue contact analyses were carried out using the g_contacts program developed by Bau and Grubmuller [39]. Given that 1H-1H NOEs are detectable up to a distance of approximatively 5-6 Å, we used a cut-off of 6 Å in the contacts analysis.

The contribution of individual residues to the total protein-protein interaction energy was computed using the MMPBSA.py [40] module available in Amber16. A total of 900 snapshots were extracted from the MD simulations of the CXCL12_2_/HMGB1 heterocomplex. Polar contributions to solvation energy were computed with the Onufriev, Bashford and Case model, setting the dielectric constant to 1 for the solute and 80 for the solvent [41]. Salt concentration was set to 0.2 M.

Nonpolar contributions to the solvation free energies were estimated by a term depending by the solvent-accessible surface area (SASA) setting γ to a value of 0.0072 kcal/mol/ Å^2^.

### 2.4 Clustering procedure

The sampled protein conformations were clustered with the g_cluster (GROMOS method) program available in the GROMACS software package (version 5.1.2) [41, 42]. After several clustering runs (Table S2) and an accurate visual inspection of the results, we verified that the application of an RMSD cutoff of 1.4 nm allowed us to discriminate different system conformations and to limit the number of singleton clusters simultaneously.

Twelve and eleven clusters were obtained for fr- and ds-HMGB1, respectively. For both systems, the centers of the first three clusters, which in both cases accounted for more than 90% of the sampled conformations, were selected for further analysis.

### 2.5 Docking procedure

The centers of the three most populated clusters derived from analysis fr- and ds-HMGB1 MD simulations were then used in docking calculations to obtain the putative structures of the CXCL12_2_/HMGB1 heterocomplex.

For CXCL12, we used the center of the most populated cluster (75.2% of the sampled structures) obtained by clustering (RMSD cutoff 3.5 Å) the simulation of 300 ns, carried out starting from the NMR structure deposited in the PDB databank with the PDBID 2KEC [43]. MD simulations were performed with the same setup and force field parameters previously used for HMGB1, adding disulfide bonds between the pairs of cysteine residues at positions 9-34 and 11-50, respectively.

Docking calculations were performed using the HADDOCK 2.2 webserver [44]. These calculations require the user to define the residues forming the binding site and, while the residues involved in the interaction between the BoxB and CXCL12 have been identified by NMR chemical shift perturbations and reported in our previous study [16], the residues forming the BoxA binding site have not yet been defined. Therefore, for BoxA we used ‘homologous’ residues obtained aligning the structures of both HMGB1 boxes (Table 2). Only the structures of the complex with the best HADDOCK scores were kept for further analysis.

**Table 2.**
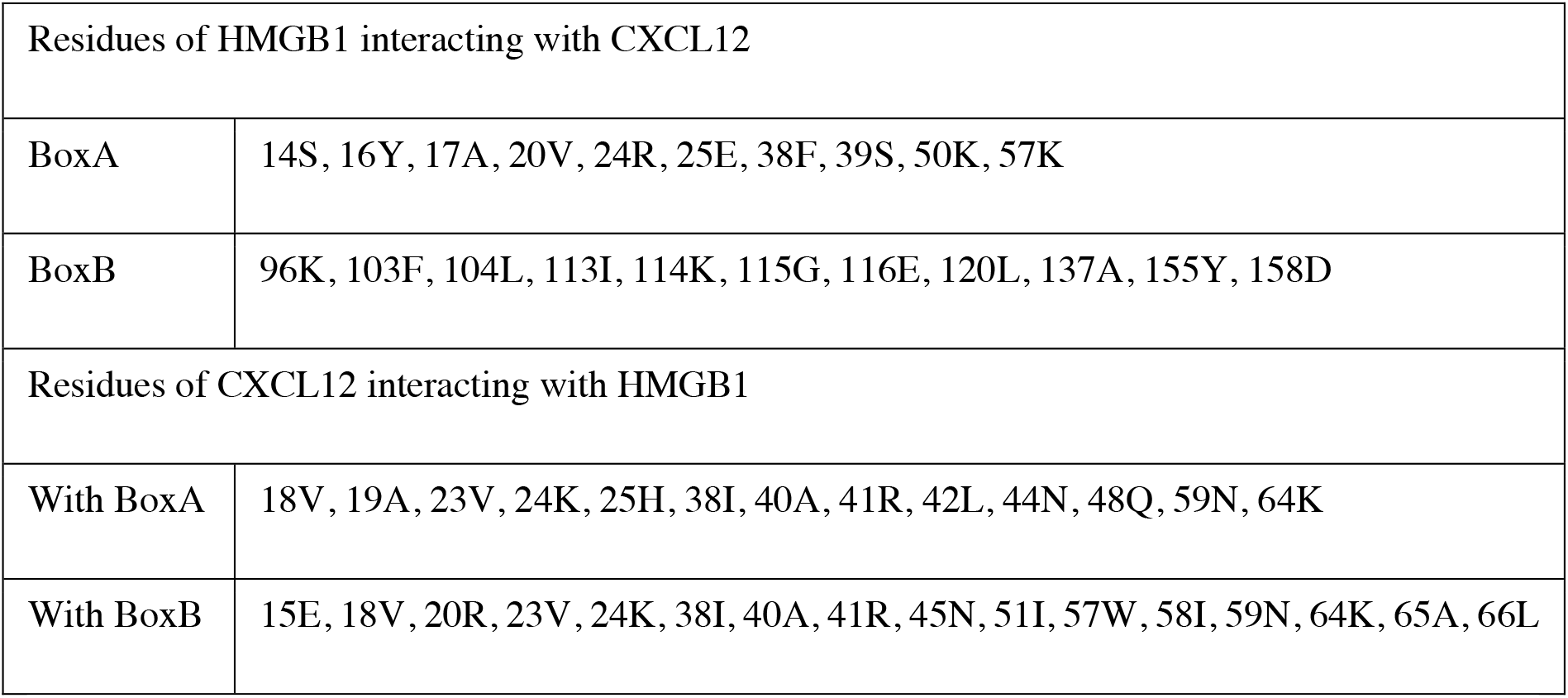
Residues involved in the interaction between HMGB1 and CXCL12 used to drive the docking procedure.

### 2.6 MD simulations of the CXCL12_2_/HMGB1 complexes

The structures of the heterocomplex obtained by docking calculations were prepared and simulated for 500 ns with the same parameters and set-up used for HMGB1 and CXCL12. During the first 200 ns, a harmonic distance restraint was applied between the centers of mass of HMGB1 and CXCL12 to optimize atomic contacts at the protein-protein interface. In particular, the force constant (k) was slowly decreased from 400 kcal/mol/Å^2^ to 0 over the first 200 ns. Then the systems were simulated for additional 300 ns. In order to increase the statistical significance of the calculations these simulations were repeated three times [45].

### 2.7 Analysis of the trajectories of the CXCL12_2_/HMGB1 complexes

The last 300 ns of the MD simulations trajectories computed for the CXCL12_2_/HMGB1 complexes were first visually analyzed to assess the stability of the complex.

Then the distance between the *N*-terminal domains of the two CXCL12 molecules were computed with the aim of determining whether the obtained CXCL12_2_/HMGB1 complexes conformations could potentially bind to and activate CXCR4 dimers.

The distance between the two binding sites in the CXCR4 receptor dimers served as the reference value. This value was determined measuring the distance between the two chemokine *N*-terminal domains (C*α* of Leu1) in the structure of a CXCR4 receptor (pdb code 4RWS [46]) in complex with a CXCL12 analog (viral macrophage inflammatory protein II (vMIP-II)).

The dimer structure was obtained applying the crystal symmetry to the deposited structure (Figure S4).

## 3 Results and discussion

### 3.1 MST investigations of the HMGB1/CXCL12 binding

Several experiments demonstrated that only fr-HMGB1 can form a heterocomplex with CXCL12 enhancing its chemotactic activity, and that CXCL12 can interacts with both BoxA and BoxB, individually [16, 18].

However, the strength of the binding between these two molecules in the two oxidation states has never been reported. Therefore, we used MST experiments to determine the dissociation constant of the heterocomplex with fr- and ds-HMGB1.

MST is a recently developed biophysical technique enabling the investigation of molecular interactions in liquid phase, i.e. without sample immobilization, measuring changes in the response to the force of a temperature gradient upon binding [22, 23].

In agreement with previously published data [17], the experiments confirmed the heterocomplex formation, with an apparent K_d_ value of 77.4 ± 16 μM (Figure 2A).Of note, using the same range of CXCL12 concentration, the heterocomplex was not detected in the presence of the ds-HMGB1 (Figure 2B), further supporting the specificity of the fr-HMGB1 for CXCL12 binding.

**Figure 2.**
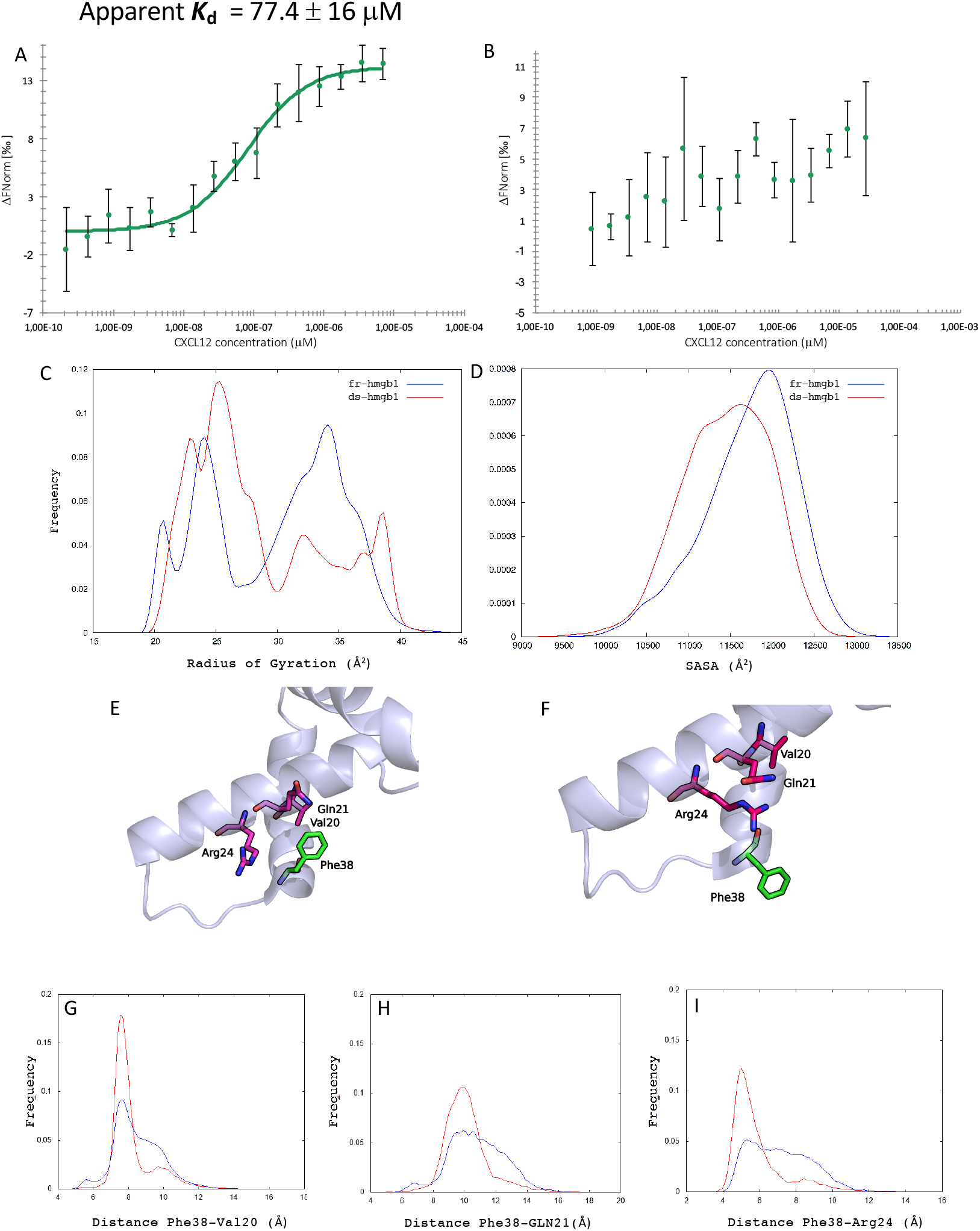
MST curve of CXCL12 titrated into labeled fr-HMGB1 (A) and ds-HMGB1 (B). (C) Histograms of the radius of gyration (RoG) and (D) solvent accessible surface area (SASA) computed using all residues of the protein. Details about the Phe38 orientation in the ds- (E, pdb code 2RTU) and fr- (F, 2YRQ) HMGB1. Histograms of the distance between the center of mass (COM) of Phe38 and COM of Val20 (G), Gln21 (H) and Arg24 (I). In all histograms, the data for fr-HMGB1 are shown in blue while those of ds-HMGB1 in red.

These findings are in line with recent data obtained by De Leo et al. [47] which report an apparent K_d_ for the CXCL12/HMGB1 in the low micromolar range.

### 3.2 HMGB1 MD simulations

According to experimental observations, only fr-HMGB1 can form a heterocomplex with CXCL12, enhancing its chemotactic activity [16, 18]. These experimental findings can be explained by different hypotheses. Indeed, the making/breaking of the disulfide bond can: (1) influence the local structure of BoxA making it unable to bind CXCL12, (2) induce a shift of the protein conformational ensemble making the HMGB1 less suitable to form the heterocomplex or, (3) the observed effect is due to a combination of the above factors. The propensity of HMGB1 to form dimers and/or tetramers has been recently shown by Helmerhorst and co-workers [48–50]. Therefore, its relevance for the different HMGB1 functions should be accurately evaluated.

Cell migration experiments [17, 20], as well as, MST measurements were performed at a fixed concentration of HMGB1 significantly below (300 nM or 10 nM) the dimerization value (K_D_) determined by SPR experiments (2 μM) [49]. Moreover, in a recent study Raggi et al. [51] determined an average concentration of HMGB1 in synovial fluids of individuals affected by oligo articular juvenile idiopathic arthritis of 2 nM. [51] From this, we can conclude that only a negligible fraction of HMGB1 is in dimeric form in the experimental conditions where the CXCL12/HMGB1 heterocomplex effect has been observed. Therefore, we simulated both the systems (fr- and ds-HMGB1) for 30 *µ*s MD considering only the monomeric form of the protein.

The simulations outputs were analyzed focusing on descriptors such as the radius of gyration (RoG, Figure 2C), the solvent accessible surface area (SASA, Figure 2D) and the RMSD with respect to NMR structure (PDB ID code 2YRQ, Figure S1), adequate to recapitulate the features of the protein conformational space.

NMR studies on ds-HMGB1, performed by Wang et al. [52] highlighted a set of 1H-1H NOE signals due to the interaction of Phe38 with Val20, Gln21, and Arg24 not detected for fr-HMGB1. As a consequence, a different orientation is assumed by Phe38 in the available HMGB1 structure (pdb codes: 2YRQ and 2RTU, Figure 2E-F).

Therefore, we focused our attention also on descriptors (distances, residue-residue, and atom-atom contacts) capable to capture the differences in the structure and dynamics of this region in the two different oxidation states (Figure 2G-I and Table S1).

RMSD analysis of BoxA (Figure S1C) resulted in very similar values for both ds-HMGB1 and fr-HMGB1, indicating that the formation of the Cys23-Cys45 disulfide bond in BoxA does not strongly alter the local conformation.

Concerning the Phe38 orientation, considering that 1H-1H NOE signals origin by short range interactions (< 5-6 Å), we monitored both the distribution of the distances between the center of mass Phe38 and the three interacting residues indicated by the NMR experiments (Val20, Gln21 and Arg24) and the percentage of the simulation time in which the atom-atom contacts responsible for the 1H-1H NOE signals are present (Table S1).

This analysis (Figure 2G, H, I and Table S1) confirmed that the presence of the disulfide bond facilitates the interaction of Phe38 with Val20, Gln21 and Arg24 however, the results of both residue-residue distance analyses and atom-atom contacts suggest that, in agreement with the dynamical nature of the system, Phe38 can flip between different conformation in both fr- and ds-HMGB1.

The RoG analysis (Figure 2C) showed a difference between the conformational spaces visited by the two systems. While two separate peaks are visible for fr-HMGB1 (the first centered at ∼24 Å and the second at ∼34 Å), only the first peak is clearly visible for ds-HMGB1. Based on this observation, the system containing the disulfide bond more frequently assumes a compact conformation than fr-HMGB1.

Finally, the SASAs for the entire protein (Figure 2D) and for BoxA and BoxB (Figure S1A-B), were estimated to evaluate the propensity of the two different HMGB1 forms to bind CXCL12. In all cases, we obtained a lower value for ds-HMGB1 than fr-HMGB1.

Summarizing, all the analyses of the simulations indicate that the presence or absence of the disulfide bond modulates the protein size and the reciprocal orientation of both the boxes and the SASA of HMGB1 without significantly altering the structure of BoxA and BoxB. As a consequence, a change in the conformational space explored by ds- or fr-HMGB1 seems to be the molecular determinant of the reduced fr-HMGB1 propensity to form a complex with two CXCL12 molecules reported in experimental studies [16–18].

### 3.2 CXCL12_2_-HMGB1 binding

To further investigate the propensity of the two HMGB1 redox states to bind CXCL12, protein-protein docking studies were performed. Representative structures were selected from the protein ensembles obtained by MD simulations by cluster analysis.

In the case of fr-HMGB1, the two most populated clusters (Figure 3A and 3C) include 55% and 20% of the conformations sampled by the system during MD simulations. Importantly, in both cluster center structures, the two CXCL12 binding sites are free (i.e., not interacting with other protein regions) and potentially able to bind CXCL12, with the N-terminal domain oriented in the same direction.

**Figure 3.**
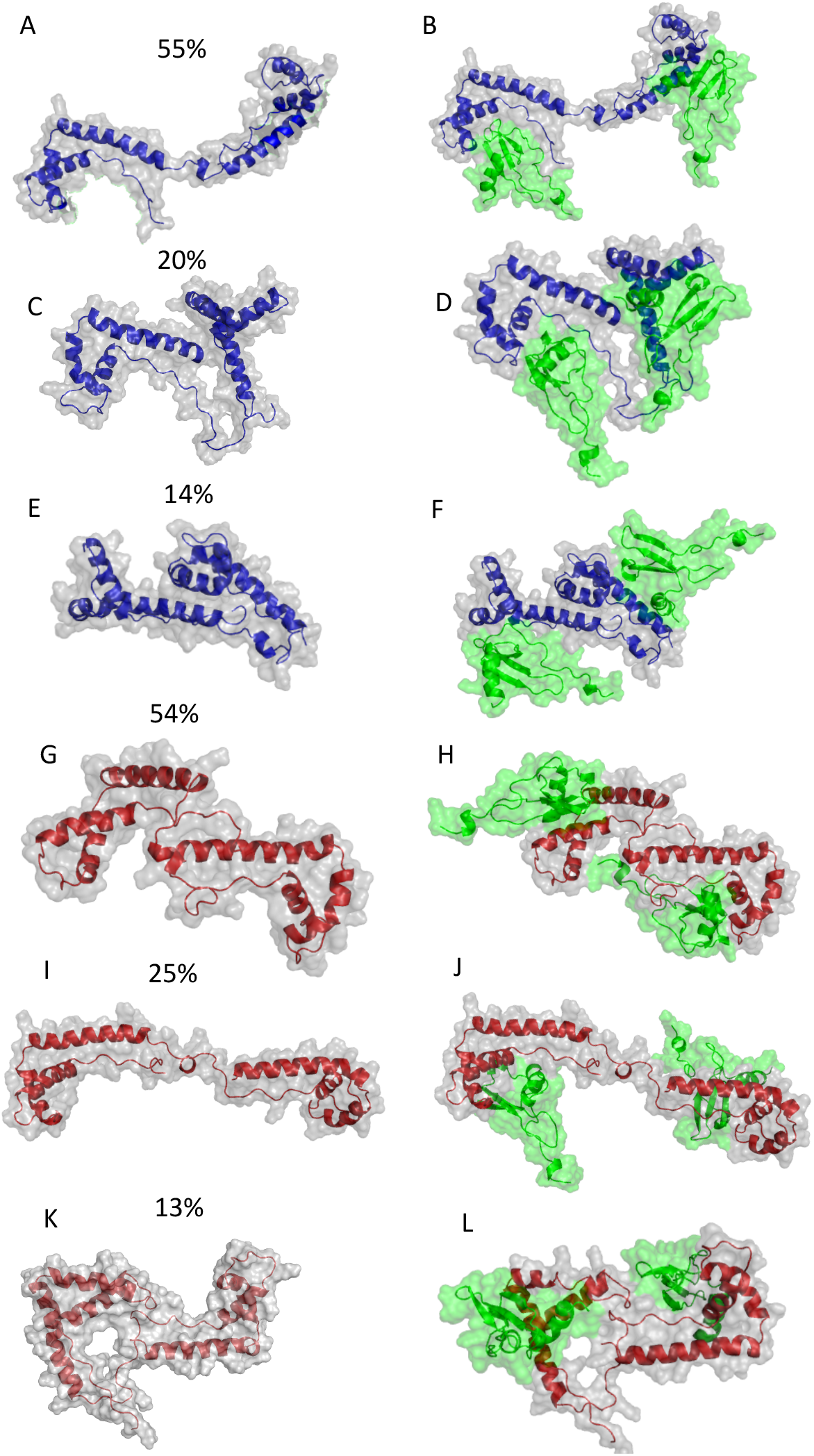
Representative conformations of the three most populated clusters (cluster centers) of fr-HMGB1 (A, C, and E) and ds-HMGB1 (G, I, and K) obtained from the cluster analysis performed using the GROMOS method.[55] The cluster size is reported as a percentage of the entire conformational ensemble. Structures of the complexes between the three most representative fr-HMGB1 (B, D, and F) and ds-HMGB1 (H, J, and L) conformations and two CXCL12 molecules (green) were obtained using protein-protein docking software HADDOCK.

**Figure 4.**
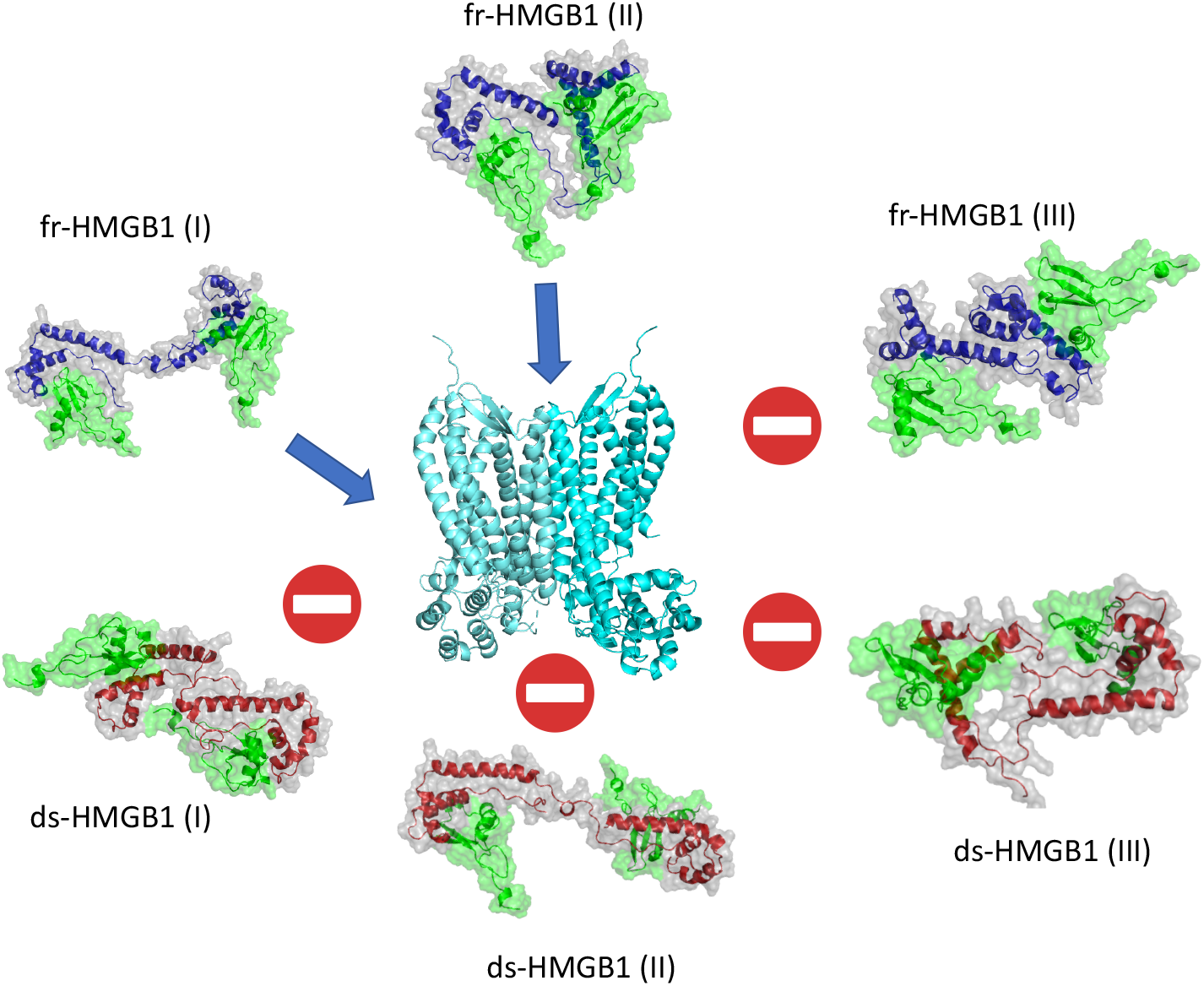
Graphical summary of the results, the blue arrow indicates that the corresponding heterocomplex can bind a CXCR4 dimer.

In contrast, the representative conformation (cluster center, Figure 3E) from the third cluster, which comprises the 14% of the generated conformational ensemble, is more compact, with the two domains interacting and, consequently, unable to bind CXCL12.

For ds-HMGB1, we observed an almost reversed trend. In this case, the first and the third most populated clusters (Figure 3G and 3K) contain 54% and 13% of the conformations, respectively. Interestingly, in both cluster centers, BoxA and BoxB are involved in reciprocal interactions that significantly limit or nullify their abilities to bind one or more CXCL12 molecules.

Only the representative conformation (center cluster) from the second cluster (Figure 3I), which accounts for 25% of the total conformations, is expanded and both domains are available to bind one CXCL12 molecule.

In summary, considering the entire conformational ensemble of fr- and ds-HMGB1 sampled during 30 *µ*s of MD simulations, we can estimate that while the ∼75% of the conformations assumed by fr-HMGB1 can activate the CXCR4 dimers, only ∼25% of the observed ds-HMGB1 conformation can do the same.

Docking calculations were performed to investigate which of the cluster centers were able to bind two CXCL12 molecules and obtain putative structures of the CXCL12/HMGB1 heterocomplexes (Figure 3B, D, F, H, J, and L). These calculations confirmed our findings from the analysis of the MD simulations trajectories. In particular, CXCL12 could be docked in the correct binding site only in the two center structures from the first two clusters from the simulations of fr-HMGB1 (fr-HMGB1(I) and fr-HMGB1(II)). Moreover, in this case, the two *N*-terminals domains of CXCL12, crucial for CXCR4 triggering [53], are oriented in the same direction, and the resulting heterocomplexes have an optimal conformation to bind a CXCR4 dimer. In contrast, the third cluster center structure fr-HMGB1 (III) is unable to bind two CXCL12 molecules due to the inaccessibility of BoxA.

In the case of ds-HMGB1, the docking of two chemokines in the correct binding site was only possible with the structure of the second cluster center. However, a visual inspection of the resulting complex (Figure 3J) reveals that the N-terminal domains of the two CXCL12 are not oriented in the same direction, making impossible the activation of CXCR4 dimers.

Docking calculations were performed using static structures, thus completely neglecting protein dynamics and the reciprocal induced fit effects. Therefore, aiming to explore the stability of the complexes obtained by docking, we simulated them for 500 ns. It should be noted that these simulations were not performed to fully explore the conformational ensemble of the complex, but to relax the system and obtain more reliable models The simulations were analyzed with a focus on the following features: (1) orientations of both binding sites for CXCL12, (2) orientations of the *N*-terminal domains of the two CXCL12 molecules and (3) stability of the complex (Table 3).

**Table 3.** Percentage of the frames sampled in the MD simulations in which the two HMGB1 domains (BoxA and BoxB) or the two N-terminal domains of CXCL12 have the same orientation.

|  |  |  |  |  |  |
| --- | --- | --- | --- | --- | --- |
| fr-HMGB1(I) |  | fr-HMGB1(II) |  | ds-HMGB1(II) |  |
| Domains | 94% | Domains | 100% | Domains | 0% |
| NT-ends | 61% | NT-ends | 92% | NT-ends | 0% |

The analysis of the MD simulations for fr-HMGB1(I) revealed that both domains are optimally oriented on the same side while the *N*-terminal domains are correctly oriented in the 61% of the analyzed conformations.

In fr-HMGB1(II) MD simulations, both domains and the *N*-terminal domains of CXCL12 were oriented in the same direction essentially for all the simulation time.

On the contrary, during the *MD* simulations of ds-HMGB1(II), which is the only conformation of the oxidized protein that can accommodate two CXCL12 molecules, both domains and the *N*-terminal of CXCL12 were oriented in opposite directions. Furthermore, the protein tended to assume conformations in which BoxA and BoxB are close to each other. Therefore, the protein conformation is more compact (Figure S3).

In order to better assess the ability of the various heterocomplexes to trigger CXCR4 dimers, we determined the optimal distance between the CXCL12 N-terminal domains (44 Å, Figure S4) analyzing the X-ray structure of the CXCR4 dimer in complex with a viral chemokine (PDB ID code 4RWS [46], see methods). This value was then compared with the average distances measured in the MD simulations (Table 4).

**Table 4.** Distance between K1 of the two CXCL12 molecules in complex with HMGB1 measured during MD simulations. The distance was only measured in simulations in which the two N-terminal domains are properly oriented to trigger CXCR4 dimers.

| Distance between K1 of CXCL12<sub>2</sub> molecules (*Ref.* = 44.0 Å) |  |  |  |
| --- | --- | --- | --- |
|  | Sim 1 | Sim 2 | Sim 3 |
| fr-HMGB1(I) | - | 44.66 ± 14.57 Å | 48.66 ± 15.03 Å |
| fr-HMGB1(II) | 53.25 ± 15.40 Å | 56.42 ± 10.81 Å | 48.88 ± 13.03 Å |
| ds-HMGB1(II) | - | - | - |

For the fr-HMGB1(I) simulations the measured average value was approximatively 44.0 Å, while for the fr-HMGB1(II) simulations, the resulting value was larger than the reference value. However, a more accurate analysis of the simulations showed that the N-terminal domains stay at the proper distance during 2/3 of the simulation time.

In summary, MD simulations performed on the complexes obtained using molecular docking lead to some interesting observations. In fact, while fr-HMGB1 forms stable heterocomplexes with the N-terminal domains of CXCL12 optimally oriented for most of the time, all complexes between CXCL12 and ds-HMGB1, sampled in our simulations, are unstable and tend to assume conformations which are not competent for the binding to CXCR4 dimers.

Lastly, aimed to determine the key interactions for the formation of fr-HMGB1(I) and (II) which emerged as potentially able to trigger a CXCR4 dimer, we computed the contribution of single residues to the protein-protein interaction energy by MM-GBSA effective binding energy decomposition (Table S4) [54].

This analysis highlighted the key role played by Phe38 in the formation of the heterocomplex with both fr-HMGB1(I) and (II) forms and indicated a weaker interaction between CXCL12 and BoxB in fr-HMGB1(II).

## 4 Conclusions

Computational studies conducted on the two redox states of HMGB1 highlighted significant differences in the conformations adopted by the fr-HMGB1 and the ds-HMGB1 forms. In particular, RoG and SASA values computed for ds-HMGB1 were significantly lower than those of fr-HMGB1, indicating that the oxidized form of HMGB1 is more compact than the reduced one, while the local structure of BoxA remained essentially unchanged over 30 μs of MD simulations.

Cluster analysis and docking calculations provided insights into the molecular determinants underlying the enhancement of CXCR4 activation induced by the heterocomplex. In fact, the analysis of these structures showed that the ∼75% of the conformations of fr-HMGB1 have BoxA and BoxB accessible for the binding of CXCL12. Furthermore, in these structures the two domains are optimally oriented to form CXCL12_2_/HMGB1 heterocomplexes competent to bind and trigger CXCR4 dimers.

In conclusion, our computational studies support the hypothesis that the absence/presence of the disulfide bond in BoxA of HMGB1, regulates the formation of CXCL12/HMGB1 heterocomplex and the enhancement of CXCR4 signaling by the modulation of the HMGB1 conformational landscape.

Furthermore, even thanking into account the intrinsic limitations of MD simulations, such as the force field accuracy, the simplified representation of the bulk and the limited conformational sampling, the results of our study provide better understanding of the CXCL12_2_/HMGB1 heterocomplex mode of action paving the way to the design of molecules capable to interfere with the CXCL12/HMGB1 heterocomplex functions.

## Abbreviations

HMGB1: High-mobility Group Box 1
fr-HMGB1: full reduced High-mobility Group Box 1
ds-HMGB1: disulfide High-mobility Group Box 1
CXCL12: C-X-C motif chemokine 12
CXCR4: C-X-C chemokine receptor type 4
TLR2 or TLR4: Toll-like Receptor 2 or 4
MD: Molecular dynamics
RMSD: root mean square deviation
SASA: solvent accessible surface area
RoG: Radius of gyration

## Supporting Information

Additional plots regarding RoG and SASA analysis; pictures of the stable compact conformations assumed by ds-HMGB1(II) with two CXCL12 molecules; representation of the CXCR4/vMIP-II complex; results of the residue-residue and atom-atom contact analysis; results of the cluster analysis carried out considering different cut-off levels, contribution of single residues to the protein-protein interaction energy determined by MMGBSA.

## Acknowledgements

AC acknowledge the Swiss National Supercomputing Center (CSCS) for the availability of high-performance computing resources. This study was supported by grants from Krebsliga Schweiz (KLS-3839-02-2016-R) and the Swiss National Science Foundation (31003A-166472 to A.C and 3100A0-143718/1 to M.U.).

