## Supplementary materials for "Oxidation state dependent conformational changes of HMGB1 regulate the formation of the CXCL12/HMGB1 heterocomplex"

**SUPPORTING INFORMATION**

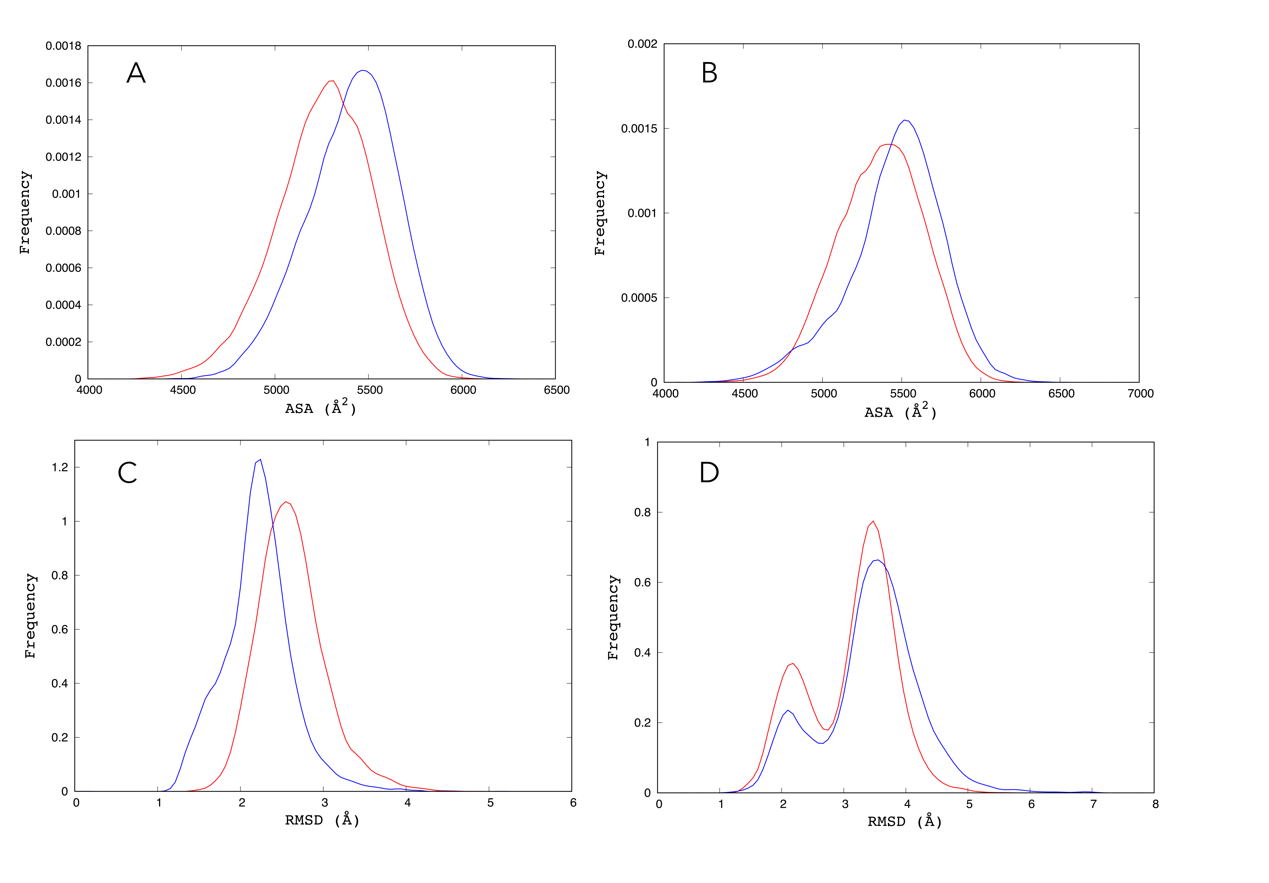

**Figure S1.** Accessible Surface Area (SASA) considering only residues from BoxA (A) and BoxB (B). RMSD of BoxA (C) (residues from Lys8 to Ile79) and BoxB (D) (residues from Lys96 to Arg163) with respect to the first structure in the NMR bundle with PDB code 2YRQ. The data are displayed in blue for fr-HMGB1 and red for ds-HMGB1.

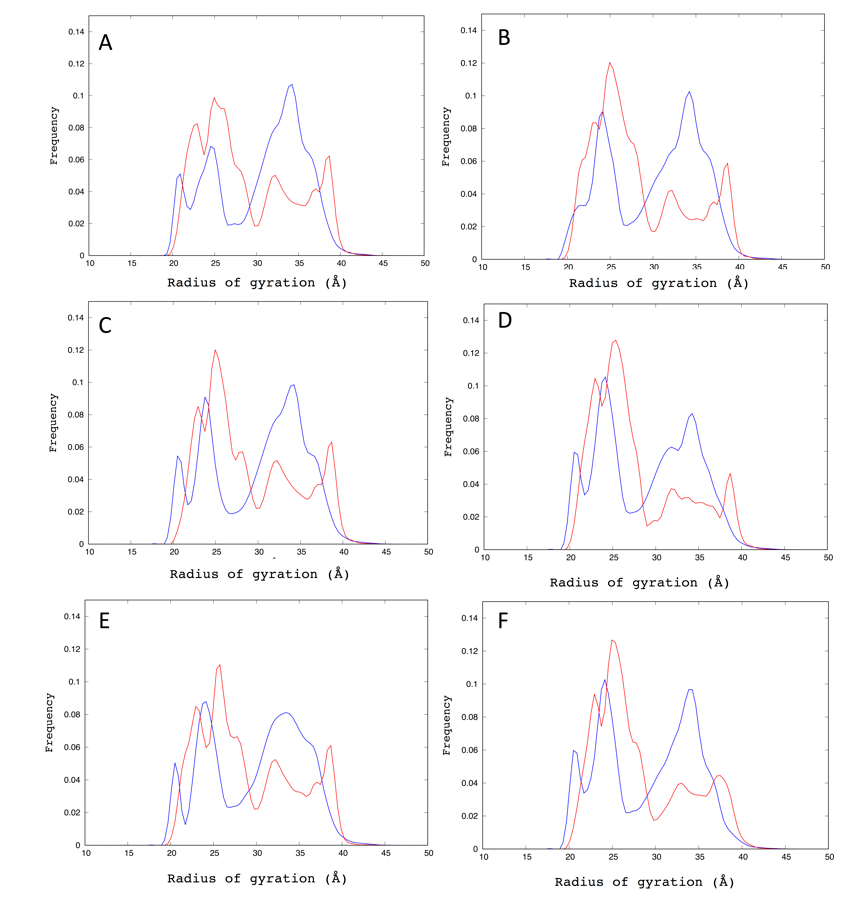

**Figure S2.** Radius of gyration (RoG) of fr-HMGB1 (blue) and ds-HMGB1 (red) of entire protein removing, each time, 12500 frames from 75000 sampled (A, B, C, D, E, F, G).

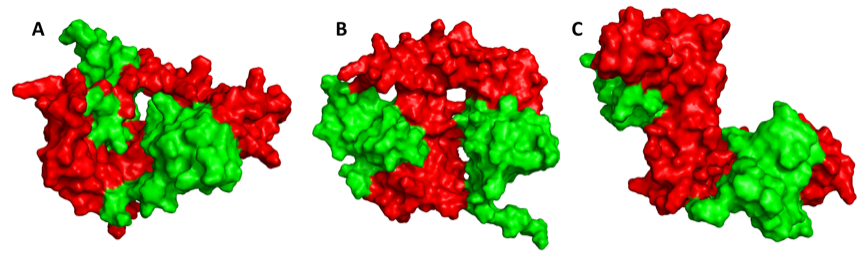
**Figure S3.** Stable compact conformations assumed by ds-HMGB1 (II), complex formed by ds-HMGB1 (red) docked with two CXCL12 molecules (green), during the first (A), the second (B) and the third (C) MD simulation.

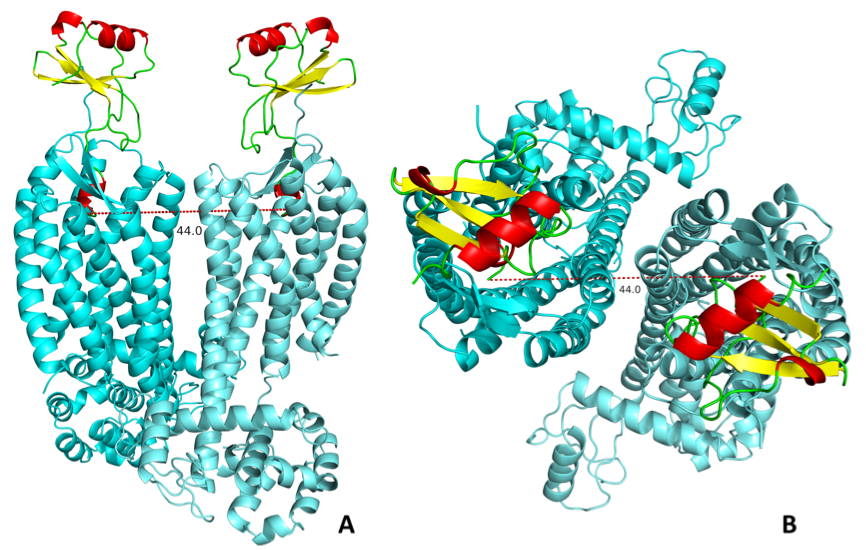
**Figure S4.** Representation of the complex CXCR4/vMIP-II (PDB ID code 4RWS) in front view (A) and top view (B) and distance between Cα of Leu1 (N-terminal tail) of the two MIP molecules (red dashes). This distance was measured to have a reference distance between binding sites of D-CXCR4 receptor.

**Table S1.**  Results of the residue-residue and atom-atom contact analysis carried out by g_contacts. In the case of hydrogen equivalent in NMR experiments an average is reported.

| Residue or atom name | Residue or atom name | Percentage of presence in fr-HMGB1 MD simulations | Percentage of presence in ds-HMGB1 MD simulations |
| --- | --- | --- | --- |
| Val20 | Phe38 | 90 | 89 |
| Gln21 | Phe38 | 55 | 76 |
| Arg24 | Phe38 | 96 | 97 |
| Val20Hβ | Phe38Hε | 27 | 33 |
| Val20Hγ1 | Phe38Hε | 62 | 61 |
| Val20Hγ1 | Phe38Hδ | 24 | 26 |
| Val20Hγ1 | Phe38Hζ | 75 | 78 |
| Arg24HN | Phe38Hδ | 15 | 35 |
| Arg24Hδ | Phe38Hδ | 39 | 56 |
| Arg24Hδ | Phe38Hζ | 56 | 39 |

**Table S2.** Results of the cluster analysis carried out considering different cut-off levels from 1 to 2 nm.

|  | **fr-HMGB1** | | | | **ds-HMGB1** | | | |
| --- | --- | --- | --- | --- | --- | --- | --- | --- |
|  | **N°** | **1°** | **2°** | **3°** | **N°** | **1°** | **2°** | **3°** |
| **1.0 nm** | 78 | 15.99% | 14.69% | 9.27% | 69 | 22.99% | 18.59% | 17.00% |
| **1.2 nm** | 33 | 30.83% | 16.46% | 13.55% | 30 | 32.73% | 24.16% | 21.79% |
| **1.4 nm** | 12 | 54.64% | 20.21% | 14.39% | 11 | 54.20% | 25.56% | 13.45% |
| **1.6 nm** | 8 | 67.00% | 23.38% | 4.98% | 7 | 72.17% | 20.22% | 5.45% |
| **1.8 nm** | 4 | 84.80% | 12.86% | 1.63% | 4 | 91.72% | 7.72% | 0.41% |
| **2.0 nm** | 4 | 99.07% | 0.89% | 0.03% | 4 | 98.42% | 1.52% | 0.05% |

**N° = Number of clusters**

**Table S3.** RMSD (Å) between all the cluster center structures calculated considering only the backbone atoms.

|  | fr-HMGB1 (I) | fr-HMGB1 (II) | fr-HMGB1 (III) | ds-HMGB1 (I) | fr-HMGB1 (II) | fr-HMGB1 (III) |
| --- | --- | --- | --- | --- | --- | --- |
| fr-HMGB1 (I) | - | 20.3 | 23.4 | 17.4 | 13.0 | 20.2 |
| fr-HMGB1 (II) | 20.3 | - | 21.1 | 12.5 | 24.2 | 16.8 |
| fr-HMGB1 (II) | 23.4 | 21.1 | - | 20.9 | 26.5 | 18.4 |
| ds-HMGB1 (I) | 17.4 | 12.5 | 20.9 | - | 17.9 | 14.2 |
| ds-HMGB1 (II) | 13 | 24.2 | 26.5 | 17.9 | - | 21.2 |
| ds-HMGB1 (III) | 20.2 | 16.8 | 18.4 | 14.2 | 21.2 | - |

**Table S4.** Contribution of single residues from fr-HMGB1(I) and fr-HMGB1(II) in complex with CXCL12 to the total protein-protein interaction energy estimated by MM-GBSA.

| fr-HMGB1(I) | | | | | |
| --- | --- | --- | --- | --- | --- |
| BoxA | | | CXCL12 | | |
| Residue | Interaction Energy Contribution (Kcal/mol) | Standard Error  (Kcal/mol) | Residue | Interaction Energy Contribution (Kcal/mol) | Standard Error  (Kcal/mol) |
| Met13 | -1.1 | 0.03 | Ala21 | -1.1 | 0.03 |
| Lys3 | -1.3 | 0.04 | His17 | -1.2 | 0.04 |
| Gly4 | -1.4 | 0.03 | Asp52 | -1.5 | 0.05 |
| Arg24 | -2.2 | 0.1 | Asn22 | -1.5 | 0.04 |
| Pro6 | -2.3 | 0.04 | Arg12 | -1.7 | 0.06 |
| Phe38 | -2.6 | 0.04 | Phe13 | -2.3 | 0.04 |
| Arg10 | -2.7 | 0.07 | Phe14 | -3.5 | 0.06 |
| fr-HMGB1(I) | | | | | |
| BoxB | | | CXCL12 | | |
| Residue | Interaction Energy Contribution (Kcal/mol) | Standard Error  (Kcal/mol) | Residue | Interaction Energy Contribution (Kcal/mol) | Standard Error  (Kcal/mol) |
| Leu104 | -1 | 0.01 | Trp57 | -1.2 | 0.02 |
| Pro99 | -1.2 | 0.01 | Asn67 | -1.2 | 0.03 |
| Glu28 | -1.6 | 0.05 | Leu26 | -1.7 | 0.02 |
| Leu63 | -1.6 | 0.02 | His25 | -2.3 | 0.06 |
| Pro23 | -3.3 | 0.02 | Tyr61 | -2.9 | 0.06 |
| Lys8 | -4.4 | 0.05 | Arg20 | -5 | 0.1 |
| fr-HMGB1(II) | | | | | |
| BoxA | | | CXCL12 | | |
| Residue | Interaction Energy Contribution (Kcal/mol) | Standard Error  (Kcal/mol) | Residue | Interaction Energy Contribution (Kcal/mol) | Standard Error  (Kcal/mol) |
| Ser14 | -1.2 | 0.03 | Leu66 | -1 | 0.03 |
| Ser39 | -1.3 | 0.04 | Trp57 | -1.2 | 0.02 |
| Lys12 | -1.5 | 0.06 | Tyr7 | -1.3 | 0.05 |
| Met13 | -1.6 | 0.04 | Val23 | -1.4 | 0.02 |
| Glu40 | -1.8 | 0.05 | Leu26 | -1.6 | 0.02 |
| Tyr16 | -1.9 | 0.03 | Pr032 | -1.7 | 0.03 |
| Asn37 | -2 | 0.03 | Ala65 | -1.7 | 0.02 |
| Phe38 | -6.1 | 0.02 | Tyr61 | -1.8 | 0.03 |
|  |  |  | Arg12 | -1.9 | 0.09 |
|  |  |  | Arg20 | -2.6 | 0.06 |
|  |  |  | His25 | -2.6 | 0.03 |
| fr-HMGB1(II) | | | | | |
| BoxB | | | CXCL12 | | |
| Residue | Interaction Energy Contribution (Kcal/mol) | Standard Error  (Kcal/mol) | Residue | Interaction Energy Contribution (Kcal/mol) | Standard Error  (Kcal/mol) |
| Ile113 | -1.1 | 0.04 | Tyr61 | -1.2 | 0.03 |
| Phe103 | -1.2 | 0.04 | Arg20 | -1.6 | 0.07 |
| Leu120 | -1.3 | 0.03 |  |  |  |
| Asp124 | -1.5 | 0.05 |  |  |  |
| His117 | -1.5 | 0.05 |  |  |  |
| Pro118 | -1.6 | 0.04 |  |  |  |
| Ser121 | -1.7 | 0.03 |  |  |  |
| Arg110 | -1.8 | 0.08 |  |  |  |
| Ile122 | -2.9 | 0.08 |  |  |  |
